# Loss of pleiotropic regulatory functions in *Tannin1*, the sorghum ortholog of Arabidopsis master regulator *TTG1*

**DOI:** 10.1101/2024.10.03.615829

**Authors:** Anthony Schuh, Geoffrey P. Morris

## Abstract

Transcriptional master regulators are often targeted to improve plant traits, but antagonistic pleiotropic effects of these regulators can hamper this approach. The Myb-bHLH-WDR (MBW) complex is a broadly-conserved transcriptional regulator affecting pigmentation, biotic stress resistance, and abiotic stress tolerance. We investigated the function of sorghum grain pigmentation regulator *Tannin1*, the ortholog of Arabidopsis pleiotropic WD40 regulator *TTG1*, to test for conserved pleiotropic regulatory effects and to better understand the evolution of the MBW complex in Poaceae. We characterized genome-wide differential expression of leaf tissue using RNA sequencing in near-isogenic lines (NILs) that contrasted wildtype *Tan1* and loss-of-function *tan1-b* alleles, under optimal temperature and chilling stress. Notably, Gene Ontology analyses revealed no pathways with differential expression between *Tan1* and *tan1-b* NILs, suggesting that, in contrast to Arabidopsis *TTG1, Tannin1* has no pleiotropic regulatory role in leaves. Further, NILs had no visible difference in anthocyanin pigmentation, and no genes with known or expected function in flavonoid synthesis were differentially expressed. Genome-wide, only 18 total genes were differentially expressed between NILs, with six of these genes located inside the NIL introgression region, an observation most parsimoniously explained by *cis*-regulatory effects unrelated to *Tannin1* regulation. Comparing our findings with known function of *TTG1* orthologs in maize, rice, and Arabidopsis, we conclude that pleiotropic regulatory function in leaf tissue was likely lost in panicoid grass evolution before the sorghum-maize split. These findings inform future molecular breeding of MBW regulated traits and highlight the benefit of subfunctionalization to relieve pleiotropic constraints.

## INTRODUCTION

Master regulators are the primary genes in a regulatory cascade (Han *et al*., 2004) and underlie many key traits for plant development and environmental response, including many traits relevant for crop improvement. Master regulators of agriculturally relevant traits include Os*WRKY71* regulating biotic stress response (Liu *et al*., 2007), CBF regulators for cold response (Savitch *et al*., 2005; Zhang *et al*., 2020), and *SELF-PRUNING* for plant architecture (Silva *et al*., 2018). However, the multiple effects of master regulators may lead to antagonistic pleiotropy (Paaby and Rockman, 2013). In Arabidopsis, the Myb-bHLH-WDR (MBW) complex is a canonical pleiotropic regulator, controlling multiple epidermal traits (seed and leaf flavonoid pigments, root hairs, leaf trichomes). It consists of three subunits, with the WD40-repeat transcriptional regulator (WDR) being shared across traits (*Transparent Testa Glabrata1*; *TTG1* in Arabidopsis), while diverse myb and bHLH transcription factors allow the complex to regulate multiple phenotypes independently (Tian and Wang, 2020). MBW function is broadly conserved across plants, along with its flavonoid regulatory function (Tian and Wang, 2020). MBW genes regulate key agronomic traits such as pigmentation, seed dormancy, grain tannins, bird resistance, fungal resistance, and trichome formation in diverse crops (Carey *et al*., 2004; Ibraheem *et al*., 2010; Wu *et al*., 2012; Xie *et al*., 2019; Yang *et al*., 2021).

Manipulation of the MBW complex has become an important target for molecular breeding (Alahakoon *et al*., 2016; Zheng *et al*., 2021; Wang *et al*., 2017), but pleiotropic effects could create undesired phenotypic tradeoffs (Benech-Arnold and Rodríguez, 2018; Gu *et al*., 2011; X., Huang *et al*., 2022). The function of *OsTTG1* in rice is similar to Arabidopsis *TTG1*, regulating trichome formation and anthocyanins throughout the plant (Yang *et al*., 2021). In maize, the MBW complex has functionally diverged, and the identified WDR subunit, *Pale Aleurone Color 1* (*PAC1)*, controls seed anthocyanins but lacks regulatory function over anthocyanins in other tissues. Differing MBW regulatory functions between rice and maize suggest evolutionary divergence occurred after the separation of the Panicoideae and Oryzoideae subfamilies c. 80 million years ago (W., Huang *et al*., 2022), which leaves the role of MBW within the Panicoideae in question, including globally important crops such as sorghum, sugarcane, and several millets. Investigation of MBW function in other panicoid grasses can be used to further resolve the evolutionary history of the MBW complex in Poaceae and better understand its role in pigmentation and stress resilience in cereals.

Sorghum (*Sorghum bicolor* L. Moench) is a globally cultivated panicoid grass crop (Monk *et al*., 2014) in which three agronomically important MBW genes have been cloned. *Tannin1* (classically, *B1*) (Wu *et al*., 2012) is a WDR, *Tannin2* (classically, *B2*) (Wu *et al*., 2019) a bHLH, and *YELLOW SEED1* (*Y1*; classically, *Y*) (Ibraheem *et al*., 2010) a myb. *Tannin1* and *Tannin2* were originally identified as regulators of grain proanthocyanidins, but these genes also colocalize with quantitative trait loci (QTL) for early season chilling tolerance (germination, emergence, seedling vigor under ∼0–10 °C), particularly *Tannin1*, which colocalizes with the largest QTL *qCT04*.*62* (Marla *et al*., 2019). *Y1* was identified as a gene controlling pericarp and leaf pigmentation and has been shown to confer resistance to fungal diseases, such as anthracnose and grain mold (Boddu *et al*., 2005; Doggett, 1970; Ibraheem *et al*., 2010; Nida *et al*., 2019). Given that the sorghum genome contains many paralogs of each of the MBW components (Morris *et al*., 2013), it is possible that they have redundant function, or they have neofunctionalized or subfunctionalized into contrasting roles (Birchler and Yang, 2022).

We investigated the conservation of the master regulatory role of *TTG1* orthologs in panicoid grasses and possible pleiotropic effects of sorghum WDR transcriptional regulator *Tannin1*, with the main goal of testing two competing hypotheses (Platt, 1964) (Fig. 1). Under the first hypothesis, the pleiotropic regulatory functions of *TTG1* would be conserved in the panicoid grass lineage leading to sorghum (*Conserved ancestral pleiotropy* hypothesis; Fig. 1A) and sorghum *Tannin1* would retain pleiotropic regulatory function in leaf tissue, including anthocyanin pigmentation (Fig. 1B). Based on colocalization of *Tannin1* with chilling tolerance QTL (Marla *et al*., 2019; Schuh *et al*., 2024) and previous reports of *TTG1*’s role in chilling response in Arabidopsis via flavonoid pigmentation (Schulz *et al*., 2016) we also considered possible pleiotropic effects of *Tannin1* on chilling tolerance (Fig. 1B). Alternatively, under the second hypothesis, these pleiotropic effects could have been lost during panicoid grass evolution (*Derived loss of pleiotropy* hypothesis; Fig. 1C) and sorghum *Tannin1* would not have a pleiotropic regulatory role in leaves (Fig. 1D). Here we used transcriptome analyses of *Tannin1* near isogenic lines, which contrast for the *Tan1* wildtype allele versus the *tan1-b* loss-of-function allele, as well as transcriptome atlas analyses from other plants, to test these hypotheses.

**Figure 1.**
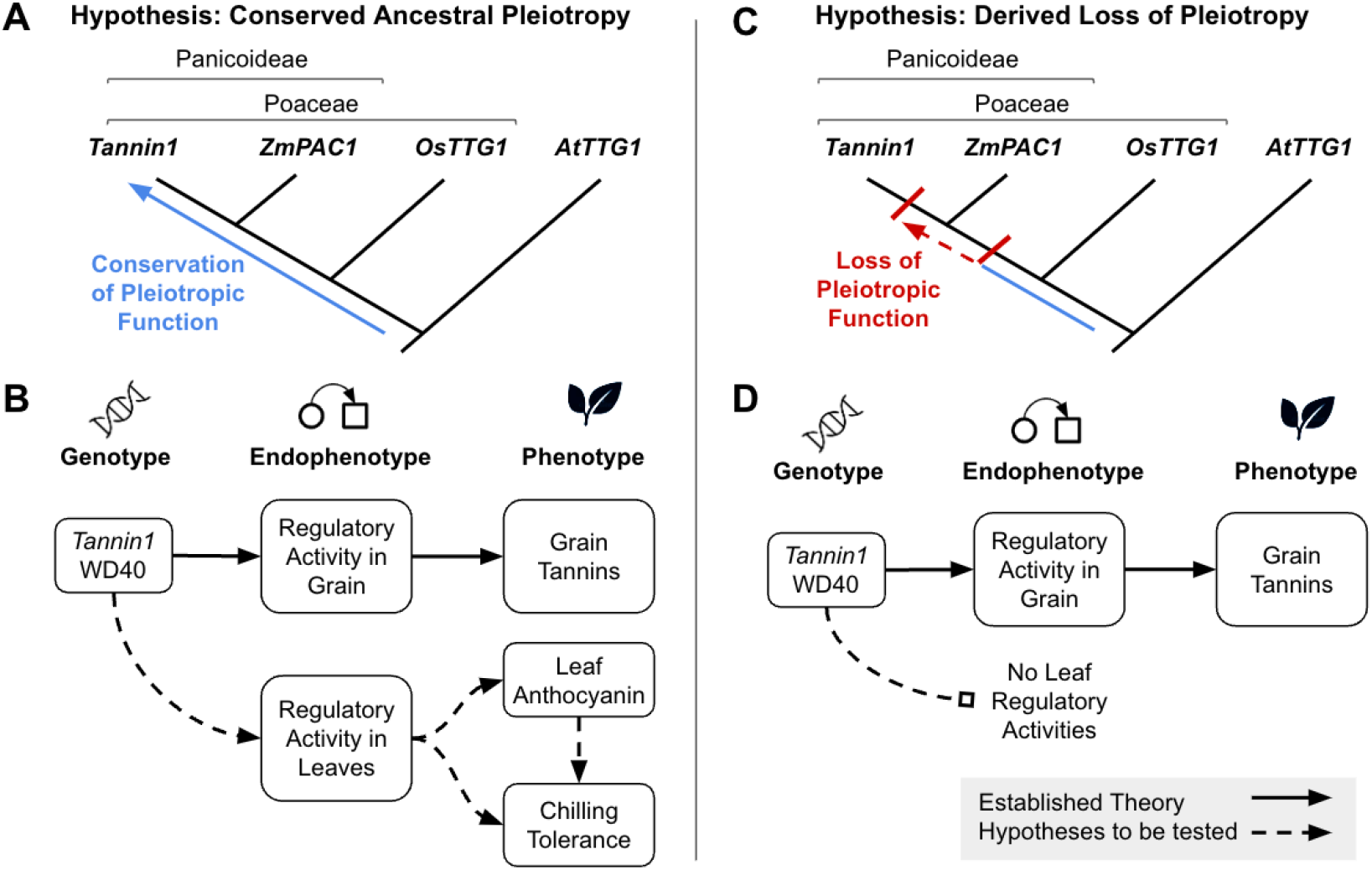
Competing hypotheses on the evolution of *TTG1* orthologs in panicoid grasses and corresponding pleiotropic functional role of the WDR protein in sorghum leaf traits. (A) Under the *conserved ancestral pleiotropy* hypothesis, *TTG1* orthologs in cereals such as sorghum, maize, and rice would have retained regulatory functions in the leaves and seed, similar to the dicot model Arabidopsis. (B) Under this hypothesis, the pleiotropic *TTG1* regulatory function in leaf would be retained in sorghum *Tannin1*, influencing the genetics of leaf anthocyanin and possibly chilling tolerance (Yang *et al*., 2021; Schulz *et al*., 2016). (C) By contrast, under the *derived loss of pleiotropy* hypothesis, pleiotropic effects of *TTG1* orthologs in leaves would have been lost during grass evolution. (D) Under this hypothesis, genetics of leaf traits would not be influenced by *TTG1* orthologs. In panel B and D, dotted lines represent hypotheses, while solid lines represent established functions (Tian and Wang, 2020).

## RESULTS

### *Tannin1* and several other WDR paralogs are widely expressed across sorghum tissues, but the other co-ortholog of *TTG1* is not

In this study, we used NILs contrasting for functional wildtype (*Tan1*) and non-functional loss-of-function (*tan1-b*) alleles under control and chilling treatments to test for a regulatory role of WDR *Tannin1* (Sobic.004G280800) in leaf tissue (Fig. 1A-B). *Tan1* NILs have a chilling-sensitive BTx623 genetic background with a 2–10 Mb introgression from chilling-tolerant kaoliang sorghum Hong Ke Zi (HKZ), including the *Tan1* allele and part of *qCT04*.*62*, notated as *pCT04*.*62+/Tan1* (Fig. 2A) (Schuh *et al*., 2024). Beyond the *Tannin1* region, *tan1-b* NILs are almost fully isogenic with BTx623 (Schuh *et al*., 2024). We first checked whether *Tannin1* is expressed in leaves, as would be expected if *Tannin1* has function in leaves (Fig. 1A). For both wildtype *Tan1* (NIL+) and *tan1-b* (NIL-), *Tannin1* is expressed in leaves, under both normal and chilled conditions, with no significant difference between the lines (Fig. 2B). Notably, however, *Tannin1* is significantly (*p* < 10^−4^) downregulated under chilling conditions in both NIL+ and NIL-genotypes (Fig. 2B). Next we considered whether Sobic.004G161600, the *Tannin1* paralog with the greatest similarity to Arabidopsis *TTG1* (73%, vs. 66% for *Tannin1*), was similarly expressed in leaves. However, there is no evidence of expression for Sobic.004G161600, in either *Tan1* or *tan1-b* NILs, in either normal or chilling conditions (Fig. 2C).

**Figure 2.**
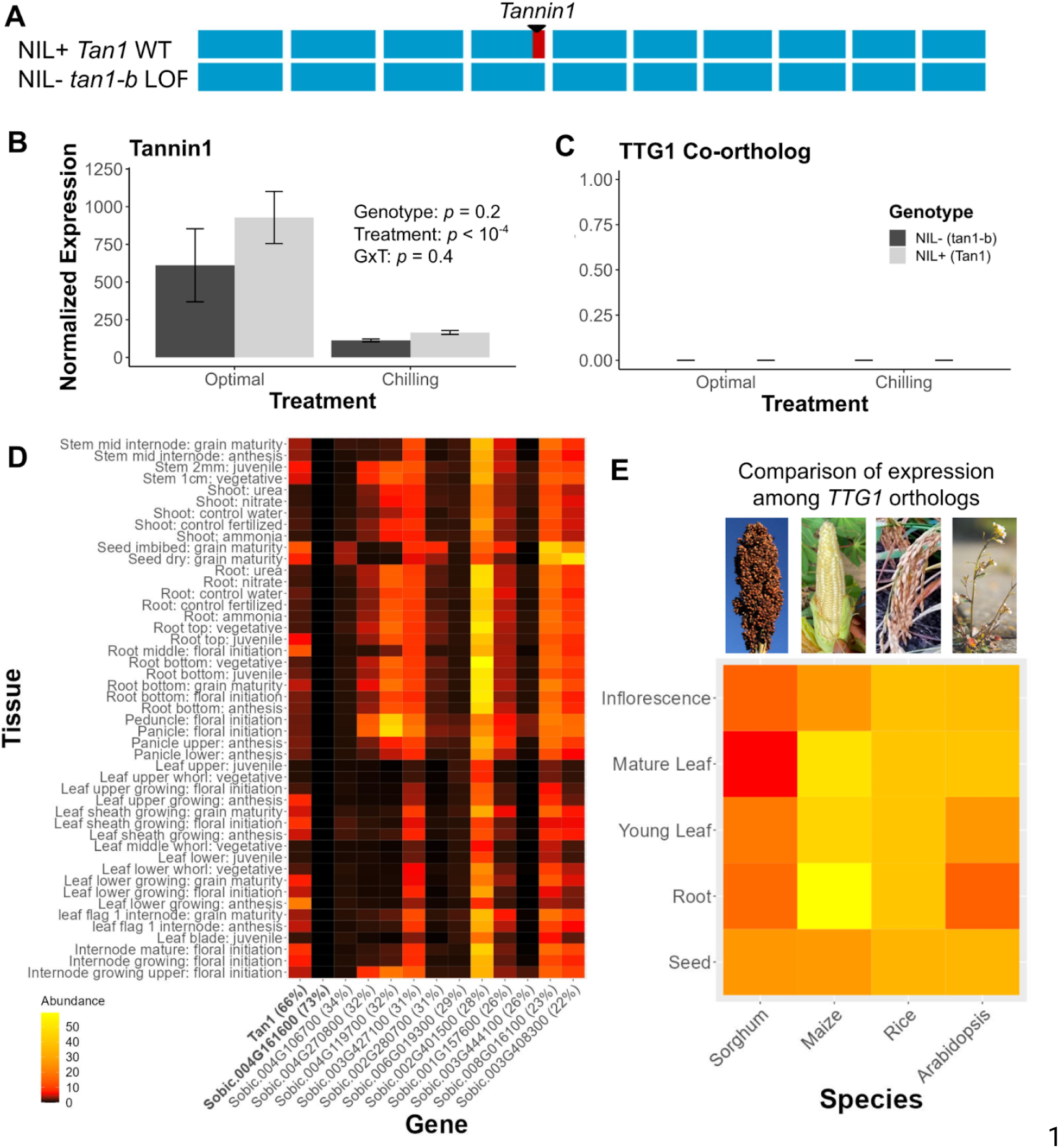
Contrasting expression patterns of *Tannin1* and other *TTG1* homologs. (A) Sliding window analysis showing inheritance of genomic regions in NILs. Blue regions are from chilling sensitive parent (BTx623) and red from chilling tolerant (HKZ). Window size is 1 Mb. (B) Expression (+/-one standard error) of *Tannin1* and (C) possible *TTG1* co-ortholog, Sobic.004G161600, in leaf tissue for NILs under control and chilling treatments (one-way ANOVA *p*-values). Expression values are the mean of six genotypic replicates and normalized using DESeq2 median of ratios. (D) Expression (FPKM) of *Tan1* and other *TTG1* homologs across sorghum tissues, from Phytozome (Goodstein *et al*., 2012). The two *TTG1* co-orthologs are indicated in bold, with percent similarity with *TTG1* in parentheses. (E) Comparison of normalized tissue expression for WDR orthologs in sorghum, maize, rice, and Arabidopsis. (Red: low relative expression, yellow: high relative expression). Tissues were manually curated by category. Data from Phytozome (sorghum), MaizeGDB (maize), BAR (rice), and TAIR (Arabidopsis) (Goodstein *et al*., 2012; Lawrence *et al*., 2004; Swarbreck *et al*., 2008; Waese and Provart, 2017). Sorghum, maize image credit: G. Morris. Rice image credit: https:/commons.wikimedia.org/wiki/File:20201102.Hengnan.Hybrid_rice_Sanyou-1.6.jpg. Arabidopsis image credit: https:/commons.wikimedia.org/wiki/File:Arabidopsis_thaliana.jpg.

To further investigate the possibility of redundant or contrasting function among WDR genes in sorghum, we characterized expression across tissues for *Tannin1* and its WDR paralogs in using publicly available data from a tissue expression atlas (Fig. 2D). *Tannin1* is not only expressed in seeds, as would be expected given its known role is grain pigmentation (Fig. 1), and in various leaf tissues, consistent with our expression analysis above; it is also widely expressed across a diverse range of tissues, including root, stem, and inflorescence (Fig. 2D). This is a similar pattern of broad expression orthologous WDR genes in maize, rice, and Arabidopsis (Fig. 2E). By contrast, there is no evidence of expression for the other sorghum *TTG1* co-ortholog, Sobic.004G161600, in any tissue or time point (Fig. 2D). Looking more broadly across paralogous WDR genes in sorghum, most of the *TTG1* homologs (with lower similarity to *TTG1*) were also not expressed in most tissues (Fig. 2D). None of the *TTG1* homologs had a leaf-specific expression profile. Notably, however, four of the paralogs (Sobic.003G427100, Sobic.002G401500, Sobic.008G016100, and Sobic.003G408300) were highly and widely expressed (including in leaf tissue at various stages), suggesting they are candidate for the WDR role in MBW function in leaves.

To further characterize *TTG1* paralogs that could play a redundant or subfunctionalized WDR role in leaf MBW regulatory complexes, we conducted a phylogenetic analysis of homologous genes from sorghum, maize, rice, and Arabidopsis (Fig. 3). The homology group from Phytozome which included *TTG1* included 71 WDR genes from the four focal species (Note, this homology group does not include all WDR genes in these species). Sorghum *Tannin1*, maize *PAC1*, and rice *OsTTG1* cluster, with Arabidopsis *TTG1* as an outgroup. Another homology group, apparently sister to the *Tannin1*-*PAC1*-*OsTTG1* group, includes the putative sorghum co-ortholog of *TTG1*, Sobic.004G161600 (red highlight) and the Arabidopsis LWD genes. Notably, the four WDR genes (other than *Tannin1*) that were highly-expressed in leaves (orange highlight; Sobic.003G427100, Sobic.002G401500, Sobic.008G016100, and Sobic.003G408300) are all relatively distantly related to the homology group that includes *TTG1, Tannin1, PAC1*, and *OsTTG1*. Among the four genes, only one Sobic.003G408300 is an ortholog of a characterized maize gene, *SHREK1* (*Shrunken and Embryo Defective Kernel 1*; v3 GRMZM2G081013; v5 Zm00001eb142980), but its function in kernel development (Liu *et al*., 2022) does not suggest a leaf WDR role for Sobic.003G408300.

**Figure 3.**
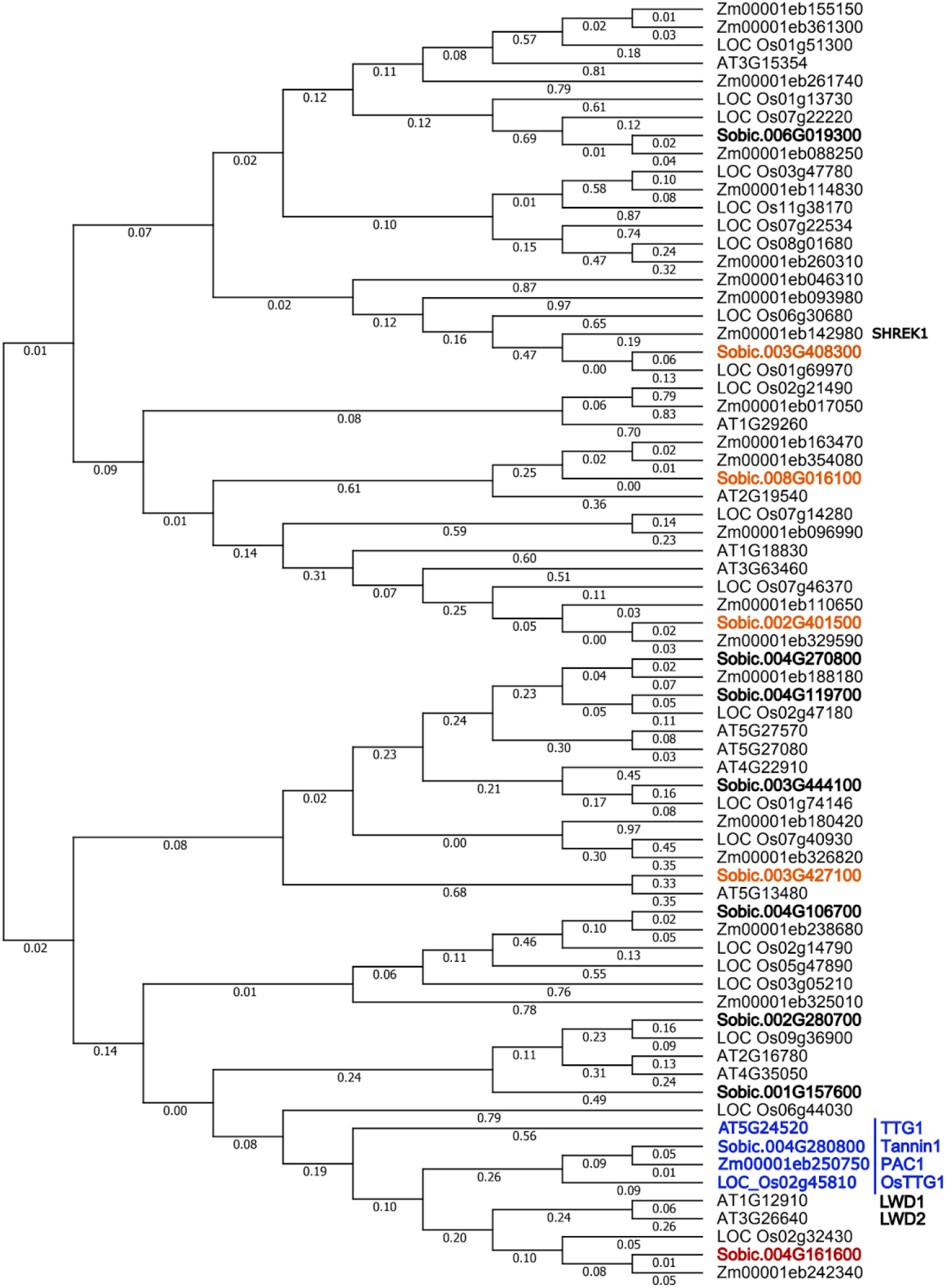
Phylogenetic analysis indicates that leaf-expressed WDR paralogs of *Tannin1* are distantly related to *Tannin1* and *TTG1*. Maximum likelihood phylogeny of WDR genes (CDS) from sorghum (v3 “Sobic” IDs), maize (v5 “Zm” IDs), rice (“LOC_OS” IDs), and Arabidopsis (“AT” IDs). Labels are branch lengths. Arabidopsis *TTG1* (At5G24520), as well as its cereal orthologs sorghum *Tannin1* (Sobic.004G280800), maize *PAC1* (Zm00001eb250750), and rice *OsTTG1* (LOC_Os02g45810), are noted in blue. Three other cloned WDR genes, maize *Shrunken and Embryo Defective Kernel (SHREK1*) and Arabidopsis *Light-regulated WD1* (*LWD1*) and *Light-regulated WD2* (*LWD2*), are also noted. The other sorghum co-ortholog of *TTG1* (Sobic.004G161600), which has no evidence of leaf expression (see Fig. 2C-D) is noted in red. Four sorghum *TTG1* homologs that were highly-expressed in leaf tissue and broadly-expressed overall (see Fig. 2D), noted in orange, are candidates for the WDR that acts in leaves.

### No *trans*-regulatory effect on *Tannin1* introgression on flavonoid biosynthesis or CBF genes in leaves

To test for evidence of pleiotropic leaf *trans*-regulatory functions of *Tannin1*, we looked for transcriptional changes in leaf tissue between NILs for genes and pathways that would be consistent with a role in leaf pigmentation or cold tolerance (Fig. 1B). If *Tannin1* has a conserved leaf pigmentation function with *TTG1* and *OsTTG1*, we would expect *trans*-regulatory effects of the *Tannin1* introgression to cause differential expression of flavonoid pathway genes between NILs. We examined all genes with a known or hypothesized (based on homology (Morris *et al*., 2013)) function in flavonoid biosynthesis and found none to be differentially expressed between the *Tannin1* NILs (Fig. 4A). Additionally, anthocyanins visibly accumulate in leaf and stem tissue for all genotypes, with no difference observed between NIL+ and NIL– genotypes (Fig. 4B).

**Figure 4.**
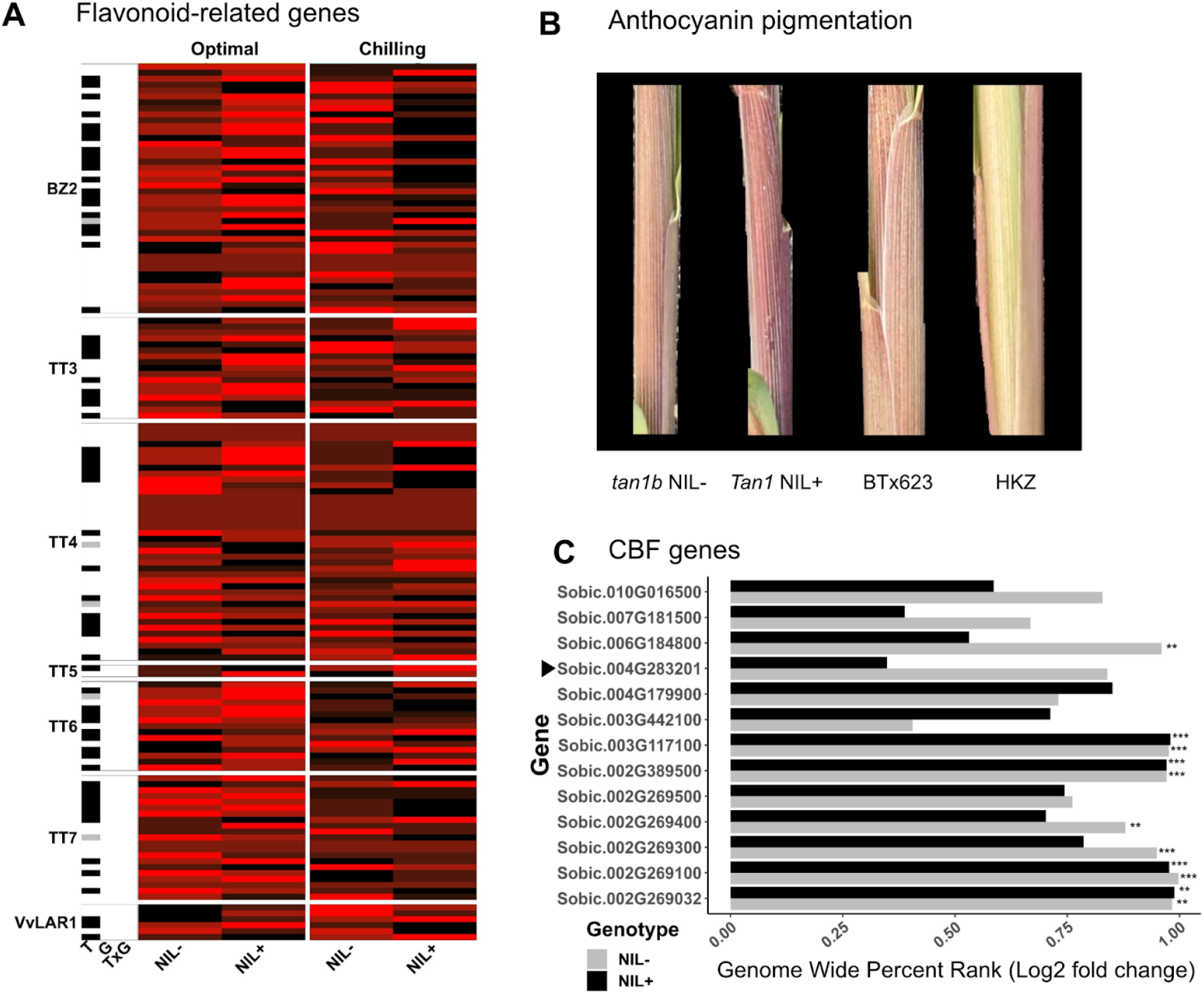
Absence of pleiotropic *trans* regulation by *Tannin1* of anthocyanins or CBF cold response regulators in leaf tissue. (A) Expression of genes with known or predicted function in flavonoid synthesis. Data is shown for genes under optimal (orange) and chilling (blue) conditions. For a given gene red indicates highest expression rank, while black represents the lowest. Median of ratios normalized mean expression from six genotypic replicates per treatment is displayed in the heatmap cell. Labels indicate the reference homolog in Arabidopsis (TT3, TT4, TT5, TT6, TT7), maize (BZ2), or grape (VvLAR) (Morris *et al*., 2013). The left sidebar shows significance of effect (black: *p* < 0.05; grey: 0.05 < *p* < 0.1; white: *p* > 0.1) for treatment (T), genotype (G), and genotype × treatment interaction (G×T). (B) Anthocyanin accumulation in stem tissue by genotype under chilling conditions. NIL parents are included, BTx623 as negative (*tan1-b*) control and HKZ as positive (*Tannin1*) control (Composite image with background set to black for clarity). (C) Genome-wide percent rank of log2 fold change in predicted CBF orthologs. Expression is measured as the mean of six genotypic replicates and normalized using DESeq2’s median of ratios method. Significance values are calculated for log2 fold change per NIL. No significant difference (*p* > 0.05) is observed for log2 fold change between NILs in any CBF ortholog. The triangle indicates the CBF homolog that is within the NIL+ introgressions. Significance codes are: * *p* < 0.05, ** *p* < 0.01, *** *p* < 0.001.

Since previous studies had posited *Tannin1* as a potential regulator of chilling tolerance (Fig. 1B) and CBF genes are known regulators of cold tolerance in Arabidopsis and maize (Thomashow, 2010), we tested whether *Tannin1* introgression had a *trans*-regulatory effect on CBF expression (Fig. 4C). Though several were up or down regulated due to chilling, no CBF orthologs were significantly differentially expressed between NILs (*p* = 1) (Fig. 4C). Further, there was no statistically significant upregulation (*p* < 0.05) for previously identified genes involved in other known chilling tolerance pathways (Marla *et al*., 2017), including lipid remodeling, NPQ, or phytohormone biosynthesis in *Tan1* versus *tan1-b* NILs (Fig. S1).

### Expression pattern suggests independent regulation among differentially expressed genes

To test for any additional *trans*-regulatory effects of the *Tannin1* introgression across the transcriptome, or *cis*-regulatory effects of other genes in the introgression, we conducted a principal component analysis (PCA; Fig. 5A). The PC1 axis, accounting for ∼42% of the total variance, separates chilling versus control treatment. The PC2 axis, accounting for 16.5% of the total expression variance, separates the Kaoliangs parent genotypes from genotypes with BTx623 genetic background (BTx623 parent line along with both NIL+ and NIL-). Thus, there is a clear structure of four discrete clusters, corresponding to genetic background by treatment combinations, starting in the top left corner and moving clockwise: Kaoliang-Chilled, Kaoliang-Control, BTx623 background-Control, and BTx623 Background-Chilled. Notably, both *Tan1* NIL+ and *tan1-b* NIL-, which have BTx623 genetic backgrounds, group with BTx623 for control and chilling treatments.

**Figure 5.**
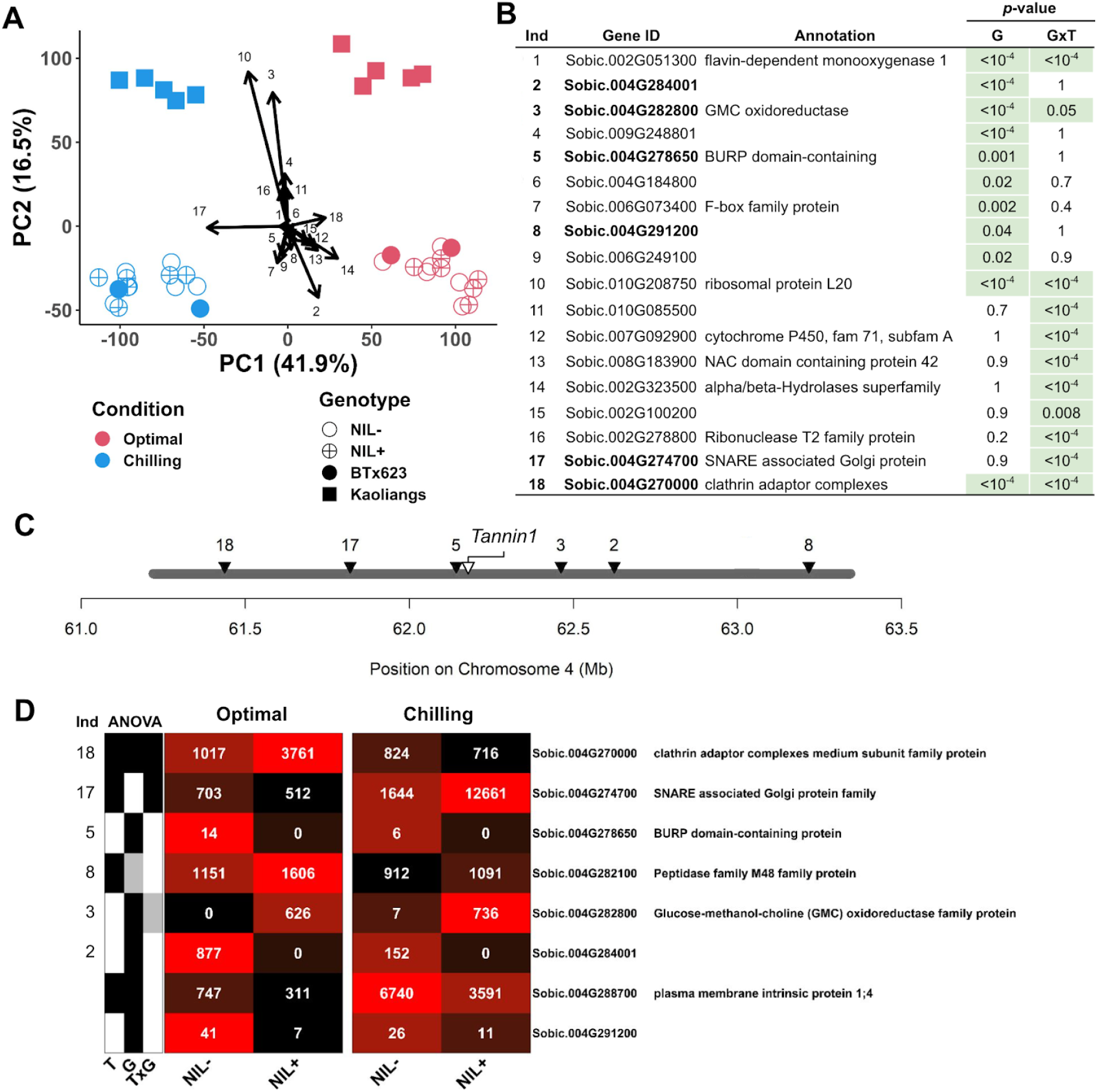
Global gene expression analysis identifies *trans* and *cis* regulatory effects of the introgression. (A) Principal coordinates analysis of global gene expression of parent lines and NILs under optimal and chilling conditions. Circles indicate BTx623 genetic background (including NILs), squares indicate kaoliangs. Filled shapes indicate parent lines while open shapes indicate NILs. PC1 axis resolves treatment while PC2 genetic background. Arrows represents principal components (scaled up 3000× for clarity) for genes with significant (*p* < 0.05; see panel B) differential expression. Direction of the arrow indicates upregulation. (B) List of differentially expressed genes (labeled with an Index, “Ind”) along with Phytozome annotations. Six differentially expressed genes located within the introgression (putatively regulated in *cis*) are indicated with bold text, while 12 other genes (putatively regulated in *trans*) are in normal text (C) Locations of differentially expressed genes with the introgression, labeled as in the previous panel. (D) Expression heatmap of introgressed genes under optimal and chilling conditions, which had significant (ANOVA *p* < 0.05) G or G×T interactions, with normalized mean expression (n = 6). Significance is on the left (black: *p* < 0.05; grey: 0.05 < *p* < 0.1; white: *p* > 0.1) and gene ID and annotation is on the right.

To identify genes that are differentially expressed due to the introgression (G effect) or due to introgression and chilling (G×T effects), we examined the expression patterns of top differentially expressed genes relative to NIL parents (Fig. 5B) and plotted these on the PCA (Fig. 5A). Overall, there is little effect of the introgression on global gene expression: among >30,000 annotated genes, just 18 genes were differentially expressed (G or G×T). Gene Ontology (GO) analysis indicated no statistical enrichment for any term (*p*-value > 0.05) and none of the genes have an annotation that suggests a role in pigmentation or chilling tolerance (Fig. 5B). Only two genes (#17: Sobic.004G274700, #1: Sobic.002G051300) have expression patterns (higher under chilling in *Tan1* NIL+) consistent with a role in induced chilling tolerance. A third of the differentially expressed genes (6/18) are in introgression itself (Fig. 5C), likely represented *cis* regulatory differences rather than *trans* regulatory effects of *Tannin1*.

Finally, to identify any other genes in the introgressed region that could contribute to phenotypic differences between the NILs via *cis* regulation (i.e. independent of *Tannin1*), we reanalyzed differential expression just within the introgression. Even with this more powerful analysis of the 225 introgressed genes, just eight (the six genes previously noted and two more) had significant (*p* < 0.05) G or G×T effects on expression (Fig. 5D). This is an enrichment within the introgression relative to a genome-wide null model (*χ*^2^ = *p* < 10^−16^). Only Sobic.004G270000 and Sobic.004G274700 exhibited G×T interactions. Only Sobic.004G274700, a putative SNARE-associated Golgi protein, is upregulated in both *Tan1* NIL+ under chilling treatment.

## DISCUSSION

### *Tannin1* is likely not a pleiotropic transcriptional regulator of leaf pigmentation or chilling response

In this study, we investigated the conservation of WDR master regulation in sorghum, using genome-wide expression patterns in *Tannin1* NILs to test for *trans*-regulatory effects that could lead to pleiotropic effects on leaf pigmentation or chilling tolerance (Fig. 1). Overall, expression patterns in NILs with contrasting *Tannin1* alleles overwhelmingly reflect BTx623 genetic background and show few regulatory differences (Fig. 4-5), which differs notably from findings in rice *OsTTG1* mutants, which had widespread transcriptome effects (Yang *et al*., 2021). The paucity of differentially expressed genes (n = 18; Fig. 5) suggests the tighter control of genetic background effects using NILs, compared to previous expression studies of chilling tolerance using diverse accessions (Marla *et al*., 2017; Chopra *et al*., 2015), allowed stronger conclusions about the genotype-phenotype relationship. There was no signal of differential regulation for leaf pigmentation (Fig. 4A-B) or chilling tolerance (Fig. 4C) pathways between *Tan1* NILs, and only 12 genes showing evidence of *trans* regulation in leaves due *Tannin1* (Fig. 5B), which would not be expected if *Tannin1* retained WDR pleiotropic regulation (Li *et al*., 2020; Yang *et al*., 2021). As far as the leaf pigmentation hypothesis (Fig. 1B), the lack of pleiotropic effects on leaf anthocyanins in *Tannin1* NILs (Fig. 4B) agrees with classical genetic studies of *Tannin1* (i.e. *B1*) and pigmentation loci (Doggett, 1970; Mace and Jordan, 2010), and further corroborates the hypothesis that *Tannin1* lacks pleiotropic leaf function (Fig. 1D).

The lack of differential regulation in chilling-associated genes between NILs (Fig. 4C) suggests that *Tannin1* has no *trans*-regulatory effect on canonical chilling tolerance pathways (Fig. 1D). This is in line with a concurrent study of morphological and ecophysiological effects of *Tannin1* introgression (Schuh *et al*., 2024), which found no developmental or physiological effects on chilling response (Fig. 4). This conclusion is surprising as both *Tannin1* and *Tannin2* are tightly co-located with chilling tolerance loci (Marla *et al*., 2019), which had suggested that grain tannins or other MBW effects contribute to chilling tolerance. It is also possible that growth chamber chilling we used does not induce the same chilling response as in the field (Marla *et al*., 2023), and we cannot entirely reject *Tannin1* as underlying *qCT04*.*62*, since other environmental stressors, such as pathogens, may drive the association (Nida *et al*., 2019). Alternatively, epistasis with other chilling tolerance alleles from Chinese sorghum (Marla *et al*., 2023; Marla *et al*., 2019), which were not introgressed here (Fig. 2A), could also account for a lack of *Tannin1* effects. If the association between *Tannin1* and chilling tolerance is due to linkage, not pleiotropy, molecular breeders could target recombinations to break the linkage (Marla *et al*., 2023) and deploy *qCT04*.*62* in early planted cropping systems (Raymundo *et al*., 2021). Further, if *qCT04*.*62* is not a pleiotropic effect of *Tannin1* it must be due to an unidentified cold tolerance gene (Schuh *et al*., 2024), which could be relevant for cold tolerance in other crops such as maize and rice.

### Relief of pleiotropic constraints due to duplication and subfunctionalization of WDR genes

The evidence that *Tannin1* does not have pleiotropic function in leaf tissues raises the question, what genes do fulfill the WDR role for other flavonoid systems in sorghum? Genetic studies (Doggett, 1970; Mace and Jordan, 2010; Ibraheem *et al*., 2010) suggest sorghum has at least four flavonoid systems, with largely independent regulatory controls: (i) proanthocyanidins (condensed tannins) in the testa (inner seed coat) (e.g. *Tannin1/B1;* WDR, *Tannin2/B2*; bHLH); (ii) anthocyanins in the seedling coleoptile (e.g. *Rs1* and *Rs2*; gene function unknown); (iii) anthocyanins in adult vegetative tissues (e.g. *P* and *Q*; gene function unknown); and (iv) phlobaphenes in the pericarp (outer seed coat) and 3-deoxyanthocyandin phytoalexins throughout the plant (e.g. *Y/Y1*; myb).

These genetic findings, along with evidence of abundant WDR duplication in the grass lineage leading to sorghum (Fig. 3), leads to the hypothesis that the WDR role has subfunctionalized, or neofunctionalized, (Birchler and Yang, 2022) across different pigmentation systems. Specifically, at least 12 other paralogous WDR regulators (Fig. 3), beyond *Tannin1*, could be considered as candidates for the WDR role for MBW function in the three other flavonoid systems listed above (ii–iv). However, the pattern of expression across tissues (Fig. 2D) and the gene phylogeny (Fig. 3), do not immediately suggest which among the WDR genes regulate other flavonoid systems (ii–iv). There is no evidence of expression of the other sorghum co-ortholog of *TTG1* (Sobic.004G161600; Fig. 2; Fig. 3), suggesting that it is either a pseudogene, or has subfunctionalized to a very limited extent of expression. Further studies of WDR paralogs in maize and sorghum (particularly the four sorghum paralogs with high leaf expression; Fig. 2-3) will be needed to identify the WDR in each system.

In rice *OsTTG1* has been shown to regulate leaf trichome development and flavonoids in several tissues, and the bHLH transcription factor Rc (orthologous to sorghum *Tannin2* and Arabidopsis *Transparent Testa 8*) regulating testa proanthocyanidins (Gu *et al*., 2011; Yang *et al*., 2021). Notably, Rc has also been shown to pleiotropically regulate seed dormancy though regulation of ABA biosynthesis (Gu *et al*., 2011), one of the findings that motivated the Conserved Ancestral Pleiotropy hypothesis (Fig. 1A-B). Taking these findings together, it is most parsimonious to infer a loss of function of leaf regulatory effects in the *PAC1*/*Tannin1* ancestor after the split from rice, but before the split between sorghum and maize (Fig. 6).

**Figure 6.**
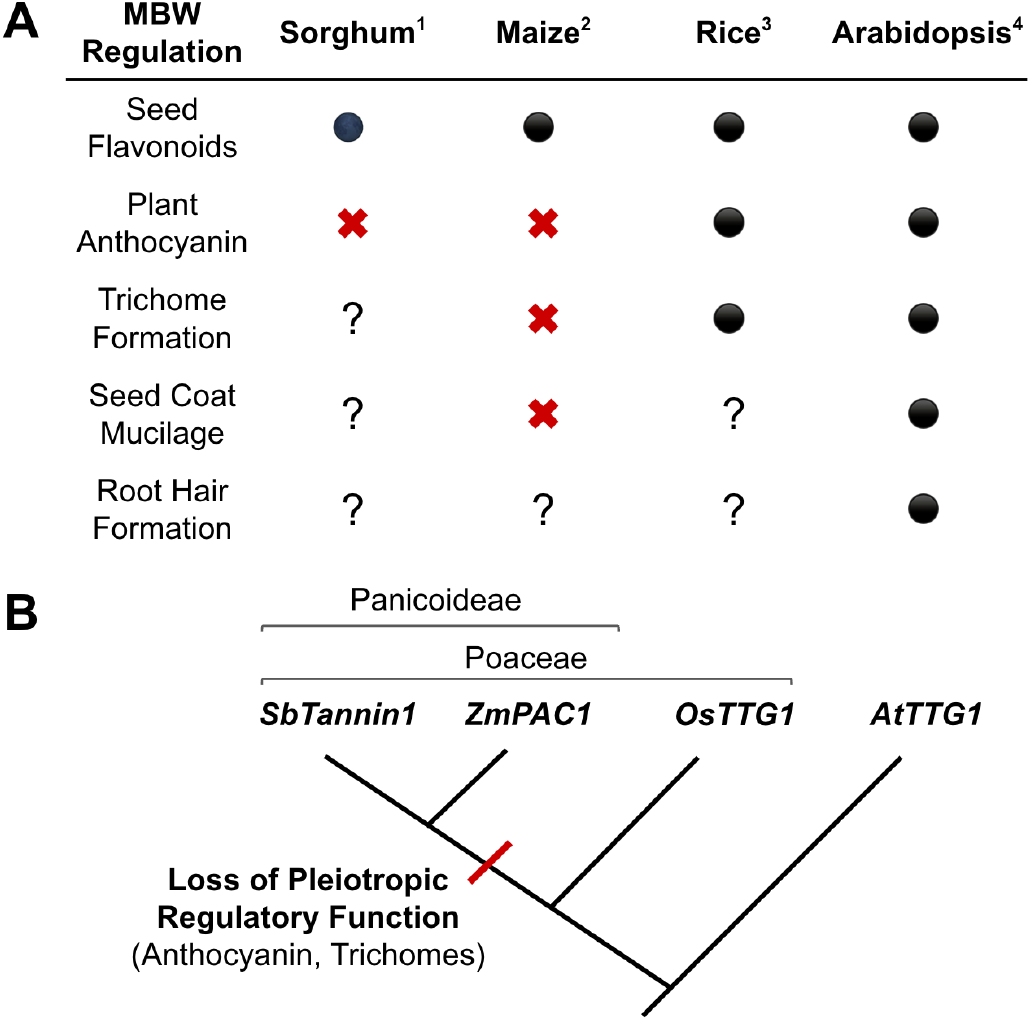
Loss of pleiotropic functions in *TTG1* orthologs in the Panicoideae after the split from rice. (A) Evidence for loss of pleiotropic function occurring before sorghum-maize speciation, but after speciation with rice. Circles indicate when MBW function is present, question mark represents unknown, and ““indicates absent function. Citations: 1, (Wu *et al*., 2012) 2, (Selinger and Chandler, 1999); 3, (Yang *et al*., 2021); 4, (Walker *et al*., 1999). (B) Hypothesis on the timing of the loss of pleiotropic functions after the panicoid grasses split from the rice lineage, but prior to the split of the maize and sorghum lineages.

While Arabidopsis *TTG1* and rice *OsTTG1* regulate flavonoids in leaves, our experiments indicate that *Tannin1* does not, which is consistent with the lack of vegetative flavonoids regulation by *PAC1* in maize (Selinger and Chandler, 1999). Thus, in maize and sorghum, it appears that multiple MBW contribute regulatory function independently in various tissues (Fig. 6) (Boddu *et al*., 2005; Ibraheem *et al*., 2010; Nida *et al*., 2019). There are two possible explanations for this independent function. Either the MBW complex has lost the need for a WDR subunit in regulating many phenotypes, and the bHLH and mybs are able to regulate independently; or, perhaps more likely, the WDR paralogs (Fig. 3) have subfunctionalized (Birchler and Yang, 2022). Our data can not fully rule out either explanation, as *Tannin1* and several other paralogs are expressed throughout the plant (Fig. 2). *Tannin1* is broadly expressed, matching the broad expression seen in *TTG1, PAC1*, and *OsTTG1* (Fig. 2) (Yang *et al*., 2021), so if *Tannin1* has subfunctionalized, it does not appear to be due to loss of expression in leaf tissue. Further, *Tannin1* and *PAC1* are both sufficient to rescue anthocyanin pigmentation in Arabidopsis *ttg1* mutants (Carey *et al*., 2004; Wu *et al*., 2012), so it is unlikely that the loss of functionality is due to coding sequence changes affecting the function of the Tannin1 or PAC1 protein. Therefore, the loss of pleiotropic function likely originates further downstream in the regulatory pathway.

## Conclusions

Overall, the hypothesis of subfunctionalization of WDRs in sorghum, and perhaps other panicoid grasses (Fig. 1C-D, 6) seems most likely, in striking contrast to the WDR master regulator model from Arabidopsis, which is the dominant model of MBW function across the plant molecular biology literature (Tian and Wang, 2020; Li *et al*., 2020; Yang *et al*., 2021). Unraveling conservation, subfunctionalized, and neofunctionalized of MBW roles will be useful for precise molecular breeding in sorghum and crops as the complex regulates multiple traits interrelatedly. The sorghum findings may also suggest hypotheses on MBW function in other related crops. For instance, several millets in the Panicoideae, understudied but globally important, have active breeding programs in developing countries (Debieu *et al*., 2017), so elucidating MBW function in sorghum and maize can inform molecular breeding in these crops. Overall, the findings illuminate the evolutionary history of the MBW complex in the Poaceae and inform strategies to improve MBW-regulated traits in cereal crops.

## MATERIAL AND METHODS

### Plant materials

The development of the *Tannin1* NILs was previously described (Schuh *et al*., 2024). Briefly, three RILs from the chilling tolerance NAM BTx623 x Hong Ke Zi (PI 567946) family were used as starting material to reduce subsequent backcrossing (Marla *et al*., 2019) and crossed to BTx623. F1 progeny were selected genotypically for heterozygosity at the QTL of interest using KASP markers and phenotypically for resemblance to BTx623, the recurrent parent. Selection and backcrossing were repeated for four generations. Four suitable BC4F1 lines were then selected and selfed. From the segregating progeny, homozygotes for both alleles of the QTL of interest were selected, making eight total BC4F2 lines. Those eight lines were then advanced to the BC4F5 generation through single seed descent generating four pairs of NIL siblings (Marla *et al*., 2023). Only NILs1-3 were used in the current study because residual heterozygosity was detected at Tannin1 locus (Schuh *et al*., 2024).

### Chilling treatment and RNA sequencing

Experiments carried out in controlled environment chambers (Conviron Model CMP6050, Manitoba, Canada) at the Plant Growth Facilities at Colorado State University in Fort Collins, CO. Experiment designs were created and randomized using a custom R v4.1.2 script (R Core Team, 2021). All plants were potted in 1.5-inch Cone-tainers using Lambert LM-HP potting soil and grown using a 12 h photoperiod and 700 μmol m^−2^ s^−1^ light intensity. After sowing 10 replicates of each genotype following the experimental design, all pots were allowed to germinate and grow under control temperature conditions, 28:25°C day:night, for approximately 14 days total. After the initial growth, half the plants were subjected to chilling conditions, 6:4°C day:night, beginning at the start of the dark photoperiod. Water was provided in excess using a bottom watering system. After a 36 hour chilling treatment, 2g of leaf tissue was collected from chilling and control plants and flash-frozen in liquid nitrogen. Frozen leaf tissue was stored at -80°C until extractions were completed. Following the manufacturer’s instructions, extractions were performed using Quick-RNA Plant Miniprep Kit (ZYMO, R2024). RNA was quantified and quality tested using a Thermo Scientific NanoDrop 2000/2000c Spectrophotometer and stored at -80°C. RNA was then sent to the Kansas State University Integrated Genomics Facility (https://www.k-state.edu/igenomics/index.html) for RT-PCR, library prep, and sequencing. cDNA was sequenced using Illumina NextSeq 500, 75 cycles, and single-read chemistry. Sequencing produced ∼2.5 GB of data per sample, or ∼30 million reads. Reads were uploaded to Illumina BaseSpace Hub by the sequencing center, and during FASTQ generation, adapter sequences were trimmed.

### Differential gene expression analysis

Reads were downloaded from Basespace Hub and mapped to BTx623 v3.1.1 reference genome (McCormick et al., 2018) using STAR v2.7.10a single pass mapping (Dobin *et al*., 2013). Subread v2.0.1 featureCounts package was then used to quantify and summarize reads (Liao *et al*., 2014). DE by genotype (G), treatment (T), and genotype by treatment (GxT) was calculated using DESeq2 v1.34.0 R package (Love *et al*., 2014). The *p*-values were obtained using the Wald test and corrected for multiple testing bias using the Benjamini-Hochberg correction. Expression was normalized across samples using DESeq2’s median of ratios method. For *cis*-regulation analysis, samples were filtered by location and significant G and GxE interactions, for other analyses, DE was examined for specific genes. Heatmaps were constructed using ComplexHeatmap v2.10.0 R package (Gu *et al*., 2016). All other plots were constructed using the ggplot2 v3.4.2 r package (Hadley Wickham, 2016). Mean expression was displayed in the heatmap cell. AgriGo: Gene Ontology Analysis Toolkit (Du *et al*., 2010) was used for Gene Ontology analysis. Data for expression analysis in other sorghum tissues were obtained from Phytozome (Goodstein *et al*., 2012). Cross-species expression comparison was performed by manually grouping tissues and assigning expression as absent, low, medium, or high for each tissue. Expression data for maize was obtained from maizegdb, rice: BAR, and Arabidopsis: TAIR (Lawrence *et al*., 2004; Swarbreck *et al*., 2008; Waese and Provart, 2017).

### Phylogenetic analyses

CDS sequences for *Tannin1* (Sobic.004G280800), *PAC1* (Zm00001eb250750), *TTG1* (AT5G24520), and *OsTTG1* (LOC_Os02g45810) were obtained from Phytozome (Goodstein *et al*., 2012). BLAST (Altschul, 1997) was used to query the Phytozome database for paralogs of WDR in their respective species.

Sequences with >25% similarity were downloaded from Phytozome and a multiple sequence alignment was created using MUSCLE (Edgar, 2004). The alignment was then trimmed using ClipKIT (Steenwyk *et al*., 2020) to maximize the accuracy of phylogenetic inference. Evolutionary analysis and tree construction were then conducted using MEGA11 (Tamura *et al*., 2021). The evolutionary relationship among genes was inferred using the Maximum Likelihood method and Kimura 2-parameter model (Kimura, 1980), and a discrete Gamma distribution was used to model evolutionary rate differences among sites. The rate variation model allowed for some sites to be evolutionarily invariable, and all positions with less than 95% site coverage were eliminated.

## Data Availability

The raw RNA sequencing data is available at the NCBI Sequence Read Archive under Bioproject accession PRJNA1168095. All other data is available at Dryad data repository under DOI 10.5061/dryad.wh70rxwxt.

## ACKNOWLEDGMENTS

This study was supported by funding from the Foundation for Food and Agriculture Research - Seeding Solution “CA18-SS-0000000094 – Bridging the Genome-to-Phenome Breeding Gap for Water-Efficient Crop Yields (G2P Bridge)” to G.P.M.

## AUTHOR CONTRIBUTIONS

TS and GPM conceived the study. TS conducted the experiments and analyzed the data. TS and GPM wrote the paper.

---

**Figure S1.**
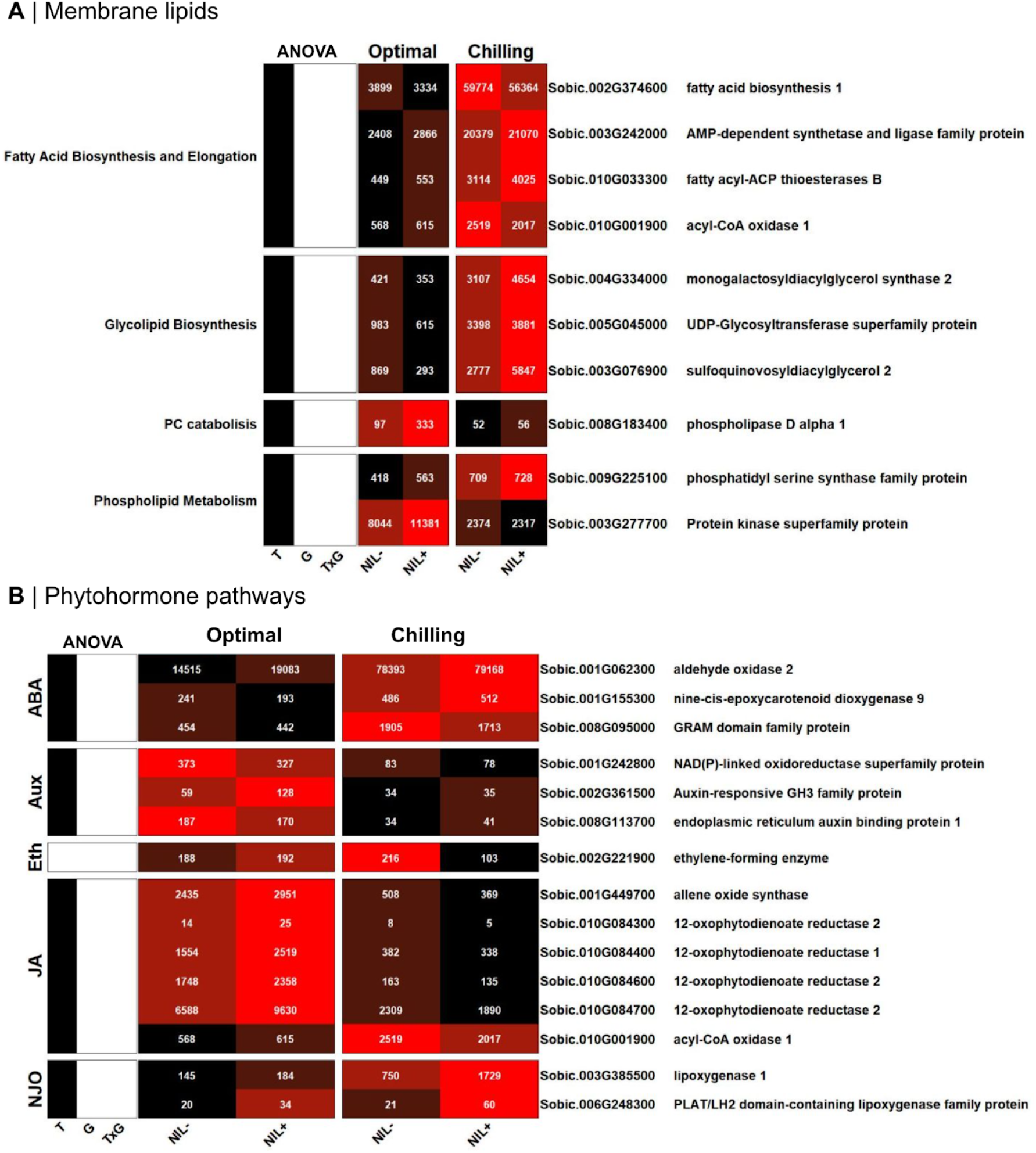
No genotype or genotype by treatment effects on expression of genes in pathways related to chilling tolerance. (A) Expression of previously identified genes with known or predicted function in lipid metabolism. Data is shown for genes under optimal and chilling conditions. For a given gene red indicates highest expression rank, while black represents the lowest. Median of ratios of normalized mean expression from six genotypic replicates per treatment is displayed in the heatmap cell. The left sidebar shows significance of effect (black: p < 0.05; grey: 0.05 < p < 0.1; white: p > 0.1) for treatment (T), genotype (G), and genotype × treatment interaction (G×T). (B) Expression of genes with known or predicted function in phytohormone biosynthesis. Leftmost labels indicate specific phytohormone, Abscisic acid, Auxin, Ethylene, Jasmonic Acid, and non-JA oxylipins.

